# Multiple Emulsions Able To Be Use For Oral Administration Of Actives Pharmaceutical Ingredients: Different Phases Physico-Chemical Parameters Study

**DOI:** 10.1101/2020.07.02.184572

**Authors:** D. L. Augustin Diaga, D. A. Rodrigue, S. Mamadou, D. S. Mouhamed, S. P. Mady, M. Gora, D. Mounibé

## Abstract

When formulating emulsions, the preliminary tests, also known as formulation tests, constitute a step which can be long and expensive because of the quantity of reagents that can be used. A rigorous methodology could thus be of great interest, that is at the aim of our study which consists of evaluating the physico-chemical parameters of different phases used to make thus multiple emulsions.

In our study, physico chemical parameters such as conductivity, pH, density, viscosity, and surface tension have been studied by direct measurement using equipment and also by means of suitable mounting.

The results showed that the pH and the surface tension have an important role in the prediction of the stability of emulsions, these latter must be of the same order of magnitude. For all phases conductivity does not have too much interest apart from helping to determine the type of the emulsion.

**STATEMENT OF SIGNIFICANCE:** Multiple emulsions are of great therapeutic interest especially in the administration of medicament which can be inactivated by digestive enzymes, moreover the researches of formulation not being often easy, a control of the physicochemical parameters of the different phases would be of great interest in rapid formulations and at low cost.

## INTRODUCTION

Emulsions are thermodynamically unstable systems, they are mixtures of two immiscible phases to be dispersed one within the other. The main goal is to keep this dispersion stable for a long time. In practice, according to Salager, “formulators of emulsions have experienced the unpleasant occurrence of the lack of reproduction of the physical properties (type, stability, viscosity) of an emulsion formulated with identical raw materials following the same rigorously definitive experimental protocol “[1, 2]. Based on this assertion, we have studied the physical parameters which allow us to find information that is relevant for obtaining emulsions with good stability and which would be carried out within a rational time.

With regard to emulsions in general, many studies have been carried out on the various techniques which improve their quality and stability. These techniques are based on compositional, formulation and process variables [1, 3-5]. Concerning the formulation and composition variables, the one which were studied are the type of surfactant, the HLB, but also the proportions of the various constituents.

## I MATERIAL

### REAGENTS

For the formulation of multiple emulsion, a mixture of Span® 80 / Tween® 80 which HLB can vary from 4.3 to 15 in used as surfactant for oil in water emulsion, the surfactant used for water in oil emulsion is Montane® 481 VG (M481VG) (HLB 4.5).

The oily or lipophilic phase used was peanut oil and the hydrophilic one consisted of Phosphate buffer Saline (PBS) pH 7.2; mixed whith Carboxymethylcellulose (CMC).

### EQUIPMENT

Equipment used consisted of :

- pH meter Schott Geräte CG820
- Conductimeter Schott Geräte CG820
- Magnetic stirrer fisher scientific
- Precision balance Ohaus explorer
- Surface tension meter Dognon-Abribat model PROLABO.

## II METHODS

For both the lipophilic and the hydrophilic phase, the physico-chemical parameters as pH, conductivity, surface tension and viscosity are studied through the above cited apparatus. To measure the viscosity, the method used consisted to form a drop of dispersed phase in the dispersing phase and in measuring the speed of migration of the drop thus formed, the device is shown in fig 1.

**Figure 1:**
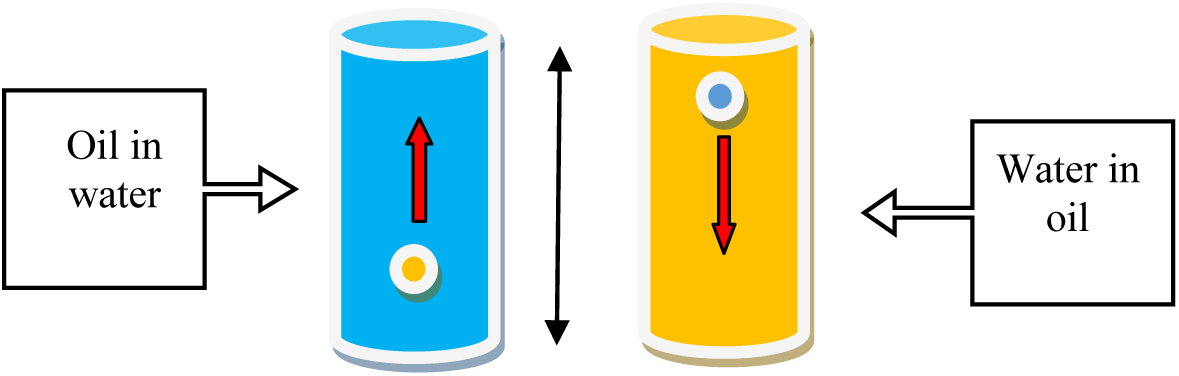
Viscosity measuring device [6].

The following relationship determines the viscosity obtained using the limiting velocity reached by a moving particle in a viscous medium :

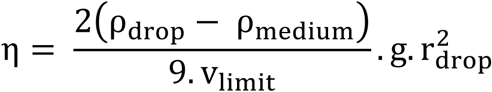

η = viscosity,ρ = density, v = velocity of the formed drop, r = radius of the formed drop, g = gravity acceleration.

It was also studied the changes of the contact angles by observing the behavior of two superposed and not mixed phases[6].

The hydrophilic phase was mixed with 1% of CMC and with the emulsifiers constituted by the span 80 / tween 80 pair at 10%. The proportion of the mixture of Span® 80 / Tween® 80 which gave different HLB are indicated in table I.

**Table I:** Proportions of span 80/Tween 80 emulsion for HLB ranging from 7 to 14.

| Span 80 | Tween 80 | BHL |
| --- | --- | --- |
| 74.77% | 25.23% | 7 |
| 65.42% | 34.58% | 8 |
| 46.73% | 53.27% | 10 |
| 28.04% | 71.96% | 12 |
| 9.35% | 90.65% | 14 |

The HLBs vary from 7 to 14 and are intended for the production of O/W emulsions.

Concerning the surfactant Montane^®^ 481 VG (M481VG) used to prepare W/O emulsion his HLB is fixe (4.5) and the proportion used are noted in table II.

**Table II:** Proportions of M481VG used for W/O emulsion

| Echantillon | Composition (%) |
| --- | --- |
| M1 | 5 |
| M2 | 6 |
| M3 | 7 |
| M4 | 8 |
| M5 | 10 |

## II RESULTS

### 1. Hydrophilic phase

The hydrophilic phase consist essentially of 1 % CMC solution for which the pH measurement gave a value of 6.8, therefore slightly acidic. For the conductivity, its measurement gave a value of 17 mS/ cm for the 1 % CMC solution and 18 mS/ cm for the 2 % solution.

The surface tension determined by the immersion blade method gave a value of 38.50 mN / m.

For the hydrophilic phase with 1% CMC mixed with the emulsifiers constituted of the span 80 / tween 80 pair at 10%, the results of the measurements of the pH, conductivity and surface tension are listed in the following table.

It is noted a decrease in pH corresponding to an increase in the degree of acidity, a slight increase in conductivity and small fluctuations in the surface tension between 29 and 31 mN / m.

### 2. Study of the lipophilic phase

The results of pH measurements of the lipophilic phase, in which the proportions of emulsifying agent vary from 0 to 8%, are shown in the table IV below.

**Table III:** Physico-chemical parameters of the CMC solution containing 10% span 80 / tween80 mixture.

| HLB | pH | Conductivity(mS/cm) | Surf. Tens.<br>(mN/m) |
| --- | --- | --- | --- |
| 7 | 5.25 | 0.91 | 30.656 |
| 8 | 5.13 | 0.96 | 31.392 |
| 10 | 4.9 | 1.01 | 29.675 |
| 12 | 4.81 | 1.02 | 30.961 |
| 14 | 4.66 | 1.15 | 31.147 |

**Table IV:** pH measurements of the lipophilic phase

| Proportions | pH |
| --- | --- |
| 0% | 2.78 |
| 2% | 3.8 |
| 4% | 3.5 |
| 6% | 4.3 |
| 8% | 4.5 |

An acidic pH is observed for all lipophilic phases, we noted an increase with the proportions of surface-active agent constituted by M481VG.

For different concentrations of montane 481 VG we also measured the surface tension which results are shown in the figure 2.

**Figure 2:**
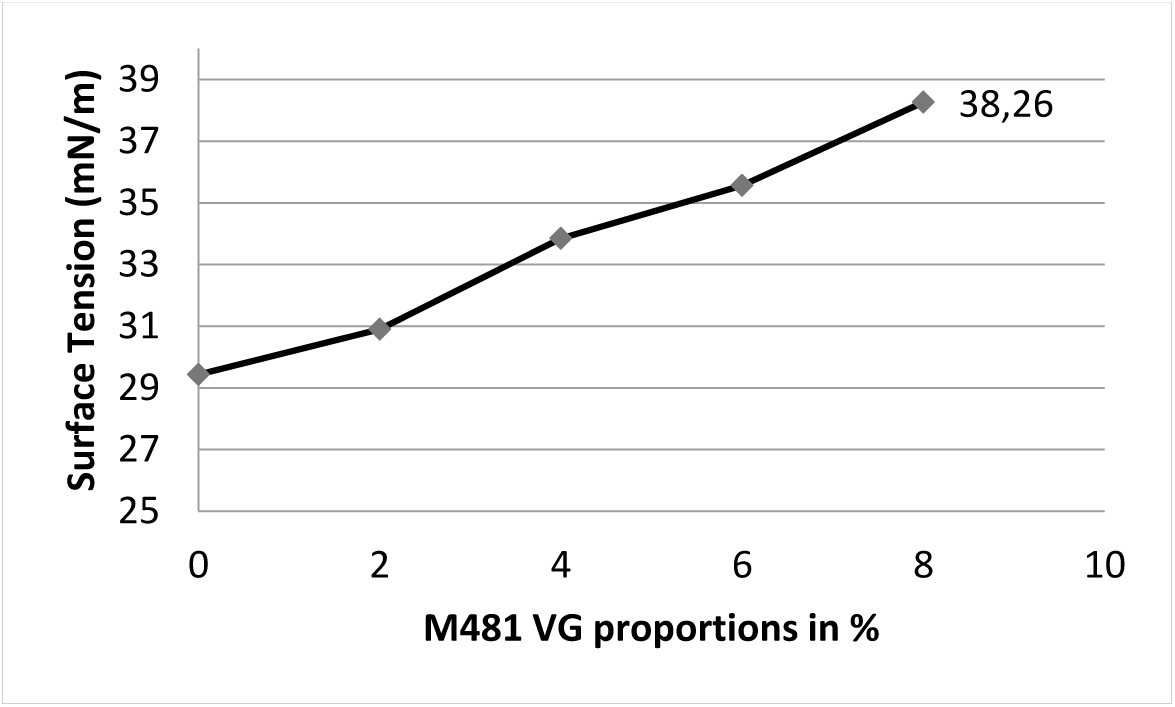
The surface tension variation curve of lipophilic phase as a function of montane 481 VG concentration

These results show that montane 481 VG increases surface tension instead of decreasing it, this is justified in so far as we have emulsions in which it is the most dense phase that must be introduced in the least dense phase, this explain the need to increase the density of the latter to avoid the sedimentation of the internal phase due to the action of gravity [7].

These changes in interfacial tension are also visible when the two solutions are superposed. A decrease in the contact angle of the oil phase with the wall of the beaker is thus observed as shown in the figure 3 following the equation F=2πRσcosθ. There is also a beginning of penetration of the oil phase into the aqueous one, which indicates a decrease in the interfacial tension.

**Figure 3:**
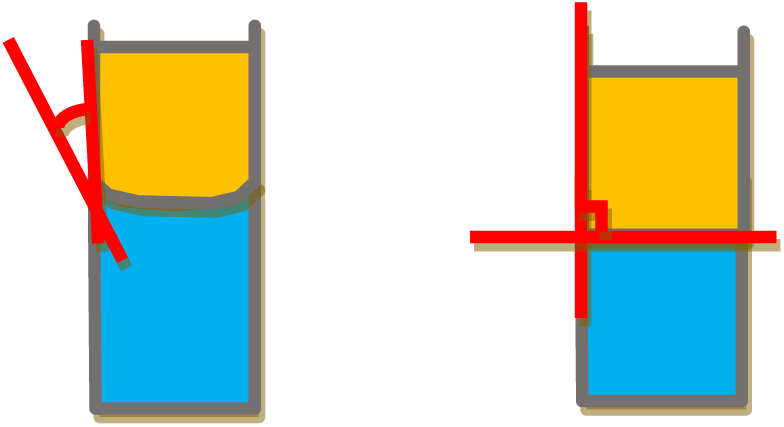
Variation of the contact angle of the lipophilic phase before (16 °) and after (90 °) addition of the surfactant [6].

For the measurement of the density, the results are given in the figure 4.

**Figure 4:**
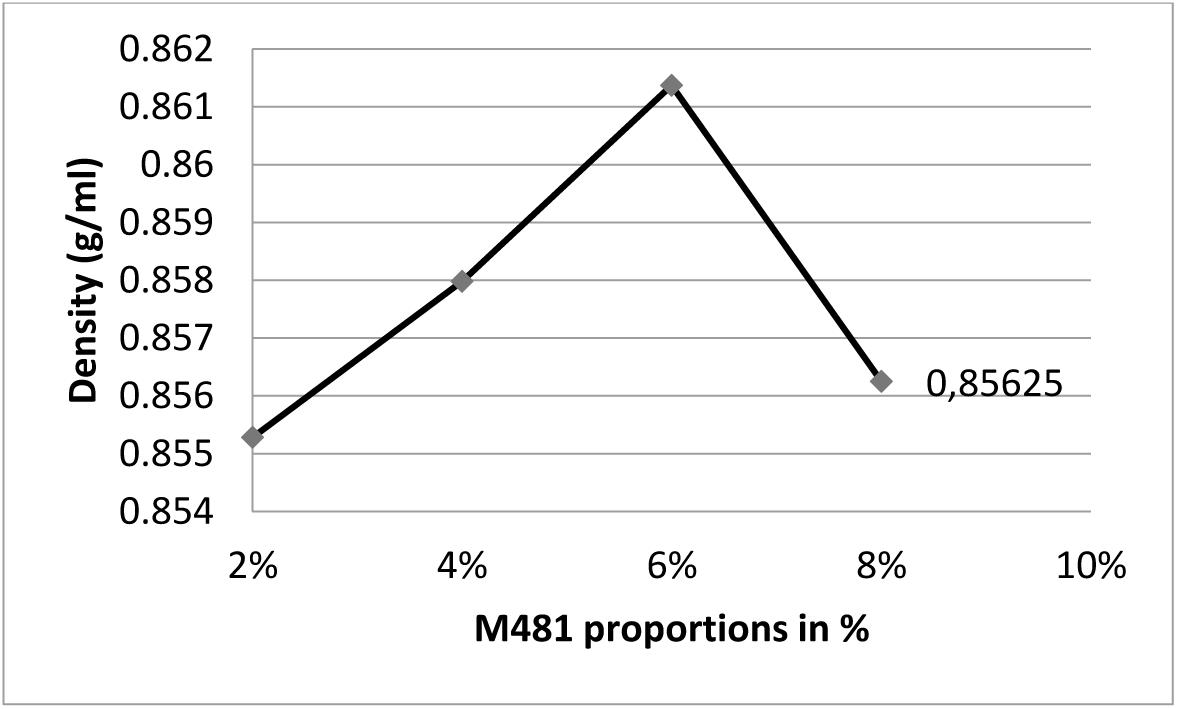
Variation of the density as function of concentration of montane 481VG

A maximal density is noted at the concentration of 6% of M481VG in the lipophilic phase.

The results of the viscosity measurements, obtained by the immersion drop method identical to the ball drop method, are shown in figure 5.

**Figure 5:**
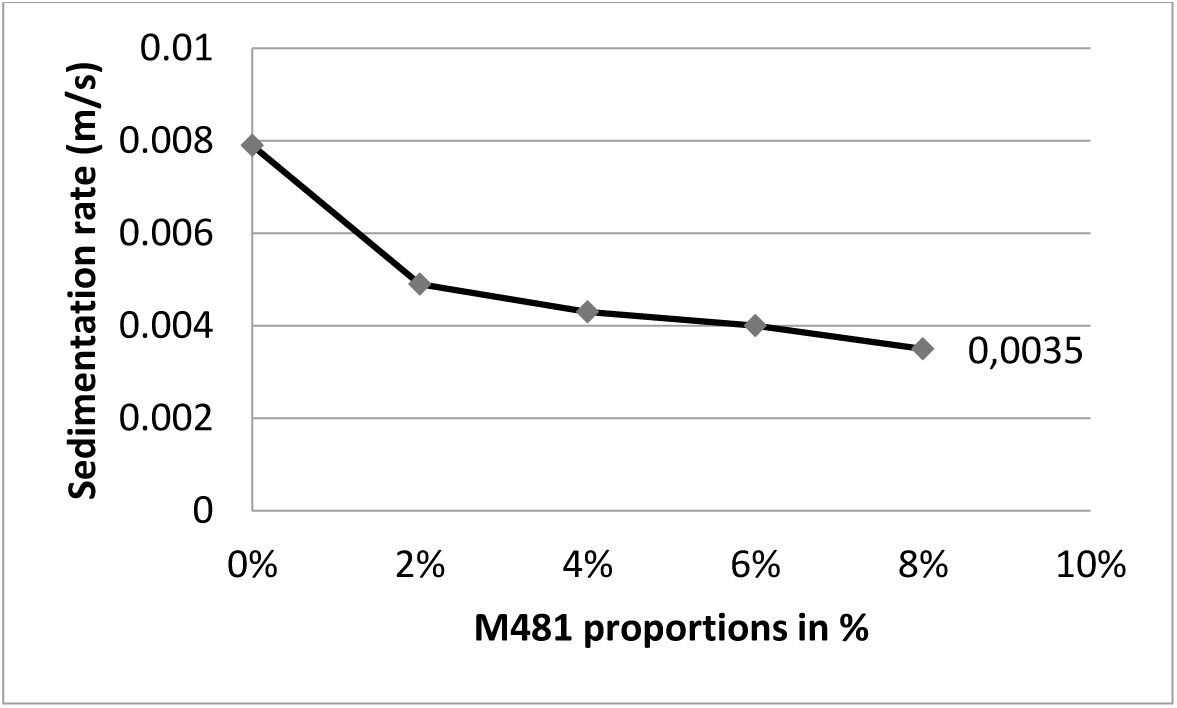
Variation of the sedimentation rate as a function of montane 481VG concentration

## III DISCUSSION

The emulsions intended to do are W/O/W emulsions type [8]. For the realization we used three types of surfactants that allowed to vary the hydrophilic / lipophilic balance. These surfactants are, tween 80 and span 80, but also montane 481VG.

Data from the literature have shown that emulsifiers for water-in-oil emulsions must have an HLB between 1 and 6, hence the use of the montane 481VG which has HLB equal to 4.5, and for oil-in-water emulsions the HLB must be between 7 and 14 hence the use of span 80 tween 80 couple [3, 9].

The study of the physico-chemical parameters such as pH, conductivity and surface tension allowed to see the variations of these properties as function of HLB especially with regard to the external hydrophilic phase. Since the goal was the formulation of multiple emulsions, the stability of the latter depends more on the external aqueous phase. For the latter, composition was of the utmost importance. Thus, as for the internal aqueous phase for which the pH is slightly acid in order of 6.8, the acidic pH is also observed for the external aqueous phase, which decreases as the HLB increases. The pH of the external hydrophilic phase to which we obtained a stable emulsion (HLB 8) of the order of 5.13 is quite close to that of the lipophilic phase with 6% emulsifier having a pH of 4.3. This acidity of the pH of the two phases is a stability indicator because an identical zeta potential can be observed around the droplets, which may be at the origin of a force of electrostatic repulsions between the droplets [10, 11]

We also noted that the HLB of span / tween 80 torque giving stable emulsions are the same as those found by ANKURMAN [12, 13].

The most interesting parameter we have studied is the surface tension at the interface of two liquids. Indeed, the main objective in the formulation of the emulsions is to reduce the interfacial tension. Therefore, if the surface tension of the aqueous phase oscillates between 29 and 30 mN/m, that of the oily phase increases in proportion to the M481VG concentration and varies from 29 to 38 mN/m. Thus, superficial tensions which observed are practically the same order of magnitude. It is also noted that the surface tension of the aqueous phase has decreased by almost half. We have tried to materialize that drop in the interfacial tension by measuring the contact angle of the aqueous phase. This shows an increase of the contact angle which has passed from 16° before addition of the surfactant to 90° after addition of 6 % of surfactant. This makes the analysis of the evolution of the surface tension of two liquids a good indicator in the prediction of stability. Besides pH and interfacial tension, the viscosity measurement can also give information about the feasibility of the emulsions. Indeed, by observing the curve of variation of the viscosity by means of the sedimentation rate of a drop of dispersed phase, it is observed that this velocity decreases and tends to stabilize around a value which,in regard to our emulsions, is 6%.

## CONCLUSION

When making emulsions, the preliminary tests, also known as formulation tests, constitute a step which can be long and expensive because of the quantity of reagents that can be used. Therefore a good methodology could be of great interest what is at the origin of our study which consists of studying the physico chemical parameters of the different phases of the emulsions to be realized. Thus the results of the study showed that the pH and the interfacial tension have an important role in the prediction of stability of emulsion, the interfacial tension of the two phases must be of the same order of magnitude. The measurement of the conductivity does not have too much interest apart from helping to determine the type of the emulsion.

## AUTHOR CONTRIBUTIONS

All authors have contributed to the research and design of this item

